# Effects of combined prenatal exposure to air pollution and maternal stress on immune and dopaminergic gene expression in the gut-brain axis

**DOI:** 10.1101/2025.09.30.678116

**Authors:** Elise M. Martin, Matthew J. Morales, Niki Y. Li, Maura C. Stoehr, Matthew J. Kern, Madeline F. Winters, Caroline J. Smith

## Abstract

Air pollution and maternal stress during pregnancy are both risk factors for neurodevelopmental disorders and often converge on the same communities. Epidemiological and animal studies suggest that maternal psychosocial stress may worsen the effects of air pollutants on neurodevelopmental outcomes. Previous work utilizing a mouse model of combined prenatal exposure to diesel exhaust particles (DEP) and maternal stress (MS) has found numerous sex-specific effects of DEP/MS exposure on neuroimmune outcomes, dopamine receptors, the gut-brain axis, and social behavior. However, it is unclear how broadly the immune landscape is shifted in the brain and intestinal epithelium following DEP/MS. Here, we analyzed immune gene expression in 5 brain regions important for social behavior and in 3 regions of the intestinal epithelium in both male and female offspring following either DEP/MS or control exposure. We found several interesting overall patterns. First, changes in expression of immune genes such as *CD11b* and *Tlr4* were concentrated in the nucleus accumbens and hippocampus. *Tlr4* and *Il-17ra* mRNA also increased in the jejunum and colon following DEP/MS, but only in females. Second, in the nucleus accumbens, catecholamine-O-methyltransferase (*Comt*) and dopamine transporter 1 (*Slc6a3*) gene expression were increased following DEP/MS indicating increased dopamine degradation at and reuptake from the synapse, respectively. Additionally, *Drd2* mRNA was decreased following DEP/MS in males. Finally, we observed numerous sex differences in immune gene expression regardless of treatment in both the brain and gut. Together, these findings suggest the nucleus accumbens is a key site for neuroimmune and dopaminergic changes following DEP/MS exposure and indicate persistent female-specific changes in intestinal immunity following these prenatal exposures.

**Highlights:**

- *Cd11b* and *Tlr4* mRNA is altered in the NAc and Hipp following DEP/MS
- Several genes indicate reduced dopamine signaling in the NAc following DEP/MS in both sexes
- Intestinal *Tlr4* and *IL-17ra* mRNA is increased in females only following DEP/MS

## 1. INTRODUCTION

Air pollution is ubiquitous in urban environments and represents a significant public health burden worldwide (Boogaard et al., 2019; B. Chen & Kan, 2008). Prenatal air pollution exposure is associated with increased risk for neurodevelopmental disorders including autism spectrum disorder (ASD; (Dutheil et al., 2021; Volk et al., 2011, 2013) and attention-deficit and hyperactivity disorder (ADHD; (Margolis et al., 2021; Min & Min, 2017; Perera et al., 2014; Yorifuji et al., 2017). Maternal psychosocial stress during the perinatal period is also associated with increased risk/severity of ASD (Alamoudi et al., 2023; Rijlaarsdam et al., 2017; Ronald et al., 2010) and other adverse neurodevelopmental consequences in children (Gutteling et al., 2006; Ronald et al., 2010; Williams et al., 2022). Moreover, air pollution disproportionately impacts under-resourced and marginalized communities that also tend to experience high levels of psychosocial stress (Jbaily et al., 2022; Rentschler & Leonova, 2023). Thus, prenatal air pollution and maternal stress exposures may converge on the same communities and synergize to increase risk for neurodevelopmental disorders. Yet, a mechanistic understanding of how these exposures combine to impact brain development remains incomplete.

Previous work in animal models has begun to address this knowledge gap with numerous studies investigating the neurodevelopmental impacts perinatal exposure to either air pollution (Chang et al., 2018; Fonken et al., 2011; Li et al., 2018; Nephew et al., 2020; Sobolewski et al., 2018; Zhou et al., 2021) or stress (H. J. Chen et al., 2024; Ehrlich & Rainnie, 2015; Gur et al., 2019). Previous work using a mouse model of *combined* prenatal diesel exhaust particle (DEP) and maternal stress (MS) exposure (DEP/MS) has observed social behavior deficits in male, but not female, offspring as compared to vehicle/control (VEH/CON; Block et al., 2022; Smith et al., 2023). DEP/MS exposure also induces maternal immune activation (MIA; (Block et al., 2022), which has been well-characterized to alter neuroimmune function and social behavior in offspring (Choi et al., 2016; Hsiao et al., 2012; Shi et al., 2003); reviewed in Zawadzka et al., 2021). In whole brain homogenates from offspring, DEP/MS exposure increases expression of toll-like receptor (TLR) 4 and the proinflammatory cytokine IL-1β while decreasing expression of the anti-inflammatory IL-10 (Bolton et al., 2013, 2017). However, these gene expression changes were not localized to specific brain regions. In the anterior cingulate cortex (ACC) and nucleus accumbens (NAc), changes have been observed in microglial structure, morphology, gene expression, and engulfment of thalamocortical synapses following DEP/MS as compared to control (Block et al., 2022; Smith et al., 2023a). However, a wide network of brain regions participates in the neural regulation of social behavior (O’Connell & Hofmann, 2011) and the effects of DEP/MS on neuroimmune gene expression have not been explored in most of them.

In addition to neuroimmune function, previous work suggests that dopaminergic signaling is impacted by prenatal exposure to DEP/MS (Smith et al., 2023a). The dopamine system is a key neural substrate on which environmental and immune challenges exert their effects (reviewed in (Filipov et al., 2002; Fone & Porkess, 2008; Kopec et al., 2019). In rats, LPS-induced MIA alters development of dopaminergic systems resulting in age-specific decreases in dopamine receptor levels in the pre-frontal cortex (Baharnoori et al., 2013). Smith et al. (2023) found that DEP/MS decreased dopamine receptor 1 (*Drd1*) and dopamine receptor 2 (*Drd2*) expression in the NAc, as well as dopaminergic input from the ventral tegmental area (VTA) to the NAc, in male but not female offspring. DA signaling in the NAc mediates social behavior, particularly behavior motivated by social reward (Gunaydin et al., 2014; Manduca et al., 2016). However, the expression of molecules related to dopamine reuptake and degradation have not been assessed in this model.

The gut microbiome is increasingly recognized to modulate neurodevelopment via the gut-brain axis (Guida et al., 2018; Lammert et al., 2018; Prehn-Kristensen et al., 2018; Wang et al., 2023) and exposure to air pollution shifts microbiome composition (Liu et al., 2021; van den Brule et al., 2021). Our previous findings show that prenatal DEP/MS shifts the composition of the gut microbiome and disrupts epithelial cytoarchitecture in offspring (Smith et al., 2023a). Moreover, cross-fostering DEP/MS exposed pups to control dams at birth rescues the social deficits associated with DEP/MS exposure in males (Smith et al., 2023a). However, it remains unclear precisely how changes to the gut impact neurodevelopment. Here we aimed to more thoroughly investigate the impact of DEP/MS exposure on immune gene expression in the gut, as cytokine signaling represents one mode by which gut microbes can communicate with the brain (Müller & Di Benedetto, 2025).

The overarching goal of this study was to expand upon previous work to characterize changes in immune and dopaminergic gene expression in a wider array of socially relevant brain regions and throughout the small and large intestine. Brain regions such as the lateral septum (LS), amygdala (AMY), and hippocampus play critical roles in social interaction, affiliation, social memory, aggression, and more(Aspesi & Choleris, 2022; Ferrara & Opendak, 2023; Menon et al., 2022). However, how they are impacted by early life experiences and immune activating events remains under explored. We collected tissue punches from 5 brain regions: the NAc, LS, AMY, and dorsal and ventral hippocampus (dHipp and vHipp, respectively); as well as from 3 segments of the intestine: the jejunum, ileum, (both small intestine) and colon (large intestine). In the brain, we assessed gene expression for *CD11b* (a microglial marker), toll-like receptor 4 (*Tlr4)*, cluster of differentiation 68 (*CD68*, a lysosomal marker), and interleukin-1β (*Il-1β*). To assess potential impacts on dopaminergic signaling, we also assessed gene expression for dopamine D2 receptor (*Drd2*), catecholamine-O-methyltransferase (*Comt*), and dopamine transporter 1 (*Slc6a3*) in the NAc, dHipp, and vHipp. We limited our investigation to these regions as these were the areas in which changes in neuroimmune signaling were observed. In the gut, we assessed the expression of *Tlr4, Il-1β, Il-17ra*, and tumor necrosis factor α (*Tnf*α). Together, these findings indicate effects of DEP/MS on immune and dopaminergic gene expression largely present in both sexes and centered in the NAc and Hipp in the brain, as well as female-specific changes in immune gene expression within the gut.

## METHODS

### 2.1 Animals

Wild-type C57Bl/6J mice were obtained from Jackson Laboratories (Bar Harbor, ME) and two generations were bred in-house before experiments commenced. Mice were group-housed with same-sex littermates in standard laboratory conditions prior to experiments (12-hour light-dark cycle, 68° F, 50% humidity, *ad libitum* food and water). Experiments were conducted in accordance with the NIH *Guide to the Care and Use of Laboratory Animals* and approved by the Institutional Animal Care and Use Committee (IACUC) of Boston College. Nulliparous females were time-mated and checked for vaginal plugs each day until a plug was observed. Presence of a vaginal plug was taken to indicate pregnancy and designated as embryonic day (E)0. Pregnant dams were pair-housed until E13, at which time they were singly housed until and after delivery.

### 2.2 DEP and VEH instillations

DEP were obtained from Dr. Staci Bilbo at Duke University and originally generated by Dr. Ian Gilmour at the Environmental Protection Agency (Durham, North Carolina). DEP or vehicle (VEH) instillations were performed as previously reported (Block et al., 2022; Bolton et al., 2013; Smith et al., 2023a). Briefly, starting on E2 and continuing every 3 days until gestation (E2, E5, E8, E11, E14, and E17; 6 instillations), dams received instillations of either 50 μL DEP (50 μg/μL) suspended in VEH (0.05% Tween20 in 1X phosphate buffered saline) or 50 μL VEH alone. During administration, dams were anesthetized with 4% isoflurane and suspended from plastic wire by their incisors. DEP or VEH was pipetted orotracheally, and inhalation was ensured by holding the tongue to prevent swallowing and covering the nasal snares for 60 seconds. Dams were then returned to their home cage and observed until they resumed typical behavior (< 5 min.).

### 2.3 Maternal stress paradigm

This model utilizes a short, prenatal version of the well-established limited nesting and bedding model (Rice et al., 2008). On E13, all dams received a cage change and normal bedding was replaced with hypoallergenic AlphaDri bedding (AlphaDri; Shepherd Specialty Papers) to reduce possible respiratory irritation. In the DEP/MS condition, an elevated aluminum mesh platform (0.4 cm x 0.9 cm mesh, McNichols Co., Tampa, FL) was placed into the cage and dams were provided 2/3 of a cotton nestlet (∼1.9g). VEH females received a full cotton nestlet and no wire mesh grate. On E17.5 (the eve of parturition) all dams were returned to CON housing conditions so that pups were reared under typical conditions. At P24, offspring were weaned and housed with same-sex littermates.

### 2.4 Tissue collection and brain punching

At P50-52, offspring were euthanized with CO_2_. To avoid litter effects, 1-4 animals per sex were obtained from 5 (VEH/CON) and 8 (DEP/MS) litters per condition. Brains were removed and flash-frozen in 2-methylbutane, then stored at -80 °C until sectioning and tissue punching. ∼10mm segments of intestinal tissue from the jejunum, ileum (proximal to the cecum), and colon (proximal to the cecum) were dissected, frozen, and stored at -80 °C until RNA extraction. Brains were mounted in a sterilized cryostat and tissue punches were collected by inserting a sterilized core sampling tool to a depth of 1 mm (Electron Microscopy Sciences). Tissue punches were collected using a 1.5 mm diameter core sampling tool. Tissue was collected bilaterally except for the LS which was on the midline. Punches were submerged in 500 µl Trizol (Thermo-Fisher Scientific) and frozen at -80 °C until RNA extraction.

### 2.5 RNA extraction

RNA was extracted using phenol-chloroform extraction. Brain punches were homogenized in 500 µl TRIzol (Thermo Fisher Scientific), and intestinal tissue was homogenized in 800 µl TRIzol. Samples were vortexed at 2000 rpm for 10 min and rested for 15 min. Chloroform (1:5 with TRIzol, Sigma Aldrich) was added and samples were vortexed at 2000 rpm for 2 min and rested for 3 min. Samples were centrifuged at 11,800 rpm for 15 min at 4° C. The aqueous phase was extracted from the phase gradient, isopropanol (1:2 with TRIzol; Sigma Aldrich) was added to the aqueous phase, and samples were centrifuged at 11,800 rpm for 10 min at 4° C. For brain punches, 2 µl Glycogen Blue (Thermo Fisher Scientific) was also added to the samples before centrifuging to improve RNA yield. RNA pellets were rinsed with ice-cold 75% ethanol (1:2 with TRIzol; Thermo Fisher Scientific) twice, centrifuging at 9000 rpm for 5 min at 4° C after ethanol was added each time. Pellets were then re-suspended in nuclease-free (NF) H_2_O (Fisher Bioreagents) and stored at -80 °C until cDNA synthesis.

### 2.6 cDNA synthesis

cDNA synthesis was performed using the QuantiTect Reverse Transcription Kit (Qiagen) according to the kit instructions. RNA quantity and quality was assessed using the NanoDrop One C (Thermo Fisher Scientific). RNA input was standardized to 1000 ng for the LS, NAc, and AMY, 800 ng for the dHipp, and 200 ng for the vHipp. A total of 12 µl suspended template RNA and NF H_2_O were combined in a ratio calculated to result in the desired amount of input RNA per sample. 2 µl gDNA wipeout buffer was added to the suspended RNA and the mixture was incubated in a Thermo Fisher Applied Biosystems MiniAmp Plus Thermal Cycler for 2 min at 42 °C, then chilled as master mix was prepared. Master mix was prepared with 1 µl reverse transcriptase (RT), 4 µl RT buffer, and 1 µl RT primer mix per sample. Master mix was added to suspended RNA. Samples were incubated for 15 min at 42 °C and 3 min at 95 °C. Samples were then diluted to 10 ng/µl and stored at -20°C unless proceeding immediately with qPCR. No-template and no-RT controls were included to verify primer specificity during qPCR.

### 2.7 Quantitative PCR (qPCR)

Quantitative PCR (qPCR) was performed using the QuantiNova SYBR Green PCR Kit (Qiagen). PCR primers were designed in-house and purchased from Integrated DNA Technologies (**Table 1**). Primers were validated to ensure that they effectively amplified the gene of interest and were specific to that gene only (i.e. melt curve analysis for multiple peaks). To prepare the SYBR master mix, 6.5 µl QuantiNova SYBR, 1 µl forward primer, 1 µl reverse primer, and 3.5 µl NF H2O per reaction were combined. 12 µl SYBR master mix per well was plated on MicroAmp Optical 96-well Reaction Plates (Applied Biosystems), then 1 µl template cDNA was added to each well. Samples were run in duplicates. The no-template and no-RT controls generated during cDNA synthesis were run on each plate to verify primer specificity. qPCR was run on a QuantStudio 3 Real-Time PCR machine (Thermo Fisher Scientific). Samples were held at 95 °C for 2 min to activate SYBR. 40 cycles of PCR were performed: samples were held at 95 °C for 5 minutes, ramped to annealing temperature at a speed of 1.6 °C/sec, and held at annealing temperature for 11 sec; then the cycle was repeated. After PCR completion a melt curve was performed.

**Table 1.**
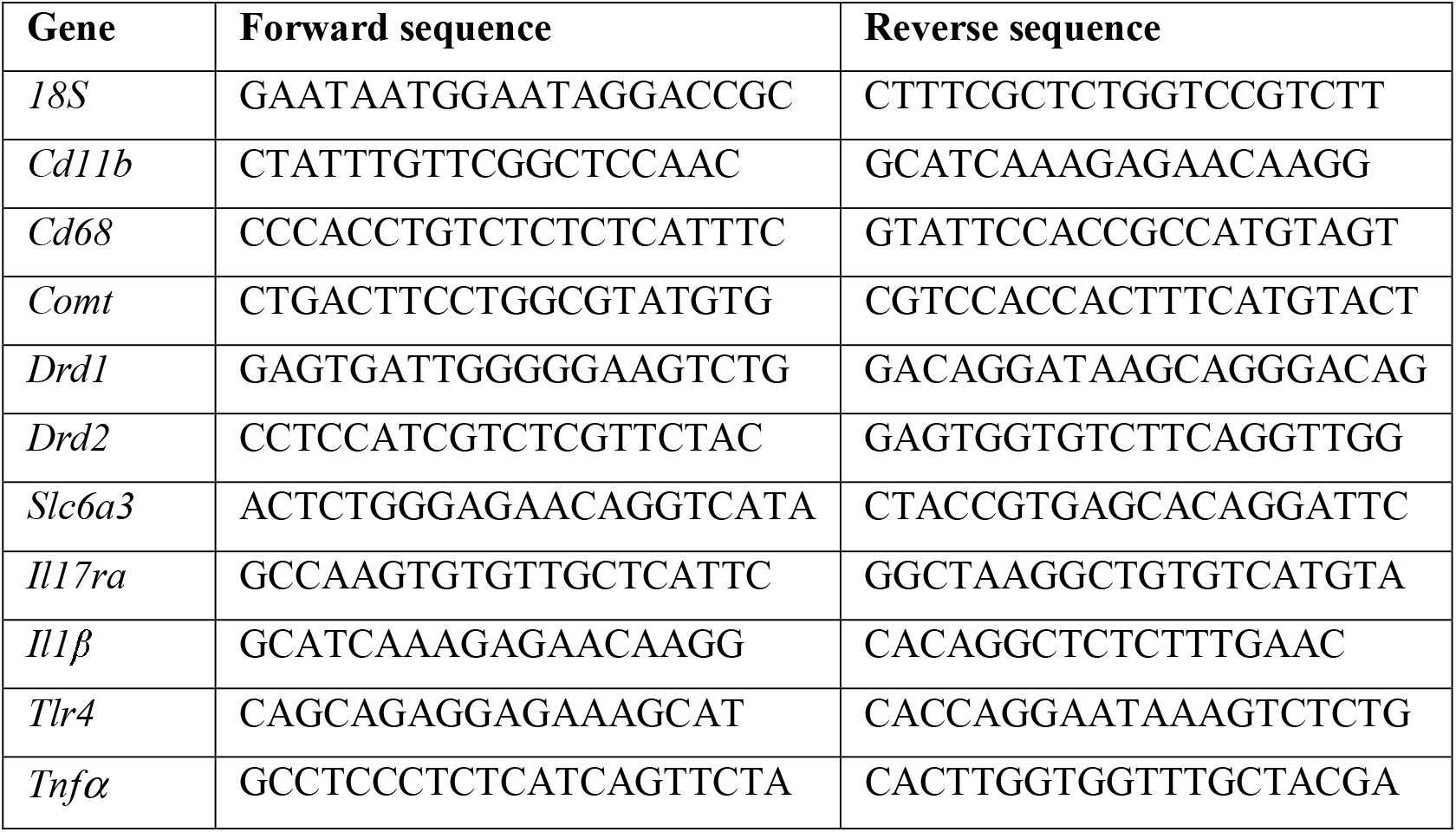
Primer sequences for quantitative PCR. Abbreviations: *18S* = 18S ribosomal RNA (housekeeping gene). *Cd11b* =cluster of differentiation 11b (integrin alpha M). *Cd68* = cluster of differentiation 68. *Comt* = catechol-O-methyltransferase. *Drd1* = dopamine receptor 1. *Drd2* = dopamine receptor 2. *Slc6a3* = dopamine transporter (DAT). *IL17ra* = interleukin-17 receptor A. *Il1β* = interleukin-1 β. *Tlr4* = toll-like receptor 4. *Tnfα* = tumor necrosis factor.

Relative gene expression was calculated using the 2-ΔΔCT method, relative to the housekeeping gene (18S) and the lowest sample on the plate (Livak & Schmittgen, 2001; Williamson et al., 2011). Microsoft Excel was used for 2-ΔΔCT calculations. If samples failed to show a melt curve with one peak or exhibited duplicate values that differed by >1 fold change they were removed from analysis before unblinding.

### 2.8 Statistical analyses

Statistical analyses were conducted using GraphPad Prism 10 software. Outliers were identified based on the ROUT outlier test (Q=1%; GraphPad prism) and removed prior to data analysis. 2-way ANOVAs [sex x treatment] were performed to assess sex, treatment, and interaction effects followed by Tukey’s *post hoc* comparisons. All data are represented as mean + SEM and statistical significance was set at p=0.05.

## RESULTS

### 3.1 DEP/MS alters the expression of immune genes in the NAc and hippocampus

We observed brain region specific effects of treatment and sex on gene expression (see **Table 2** for complete F statistics). In the NAc, DEP/MS increased *Cd11b* mRNA (**Fig. 2a**; treatment: p=0.048) and *Tlr4* mRNA expression (**Fig. 2a**; treatment: p=0.04). There were no effects of treatment or sex on *Cd68* or *Il1*β mRNA expression (**Fig. 2a**). DEP/MS also increased *Tlr4* mRNA expression in the dHipp (**Fig. 2b**; treatment p=0.03) as compared to VEH/CON, but had no effect on *Cd11b, Cd68* or *Il1*β mRNA. Females trended towards expressing more *Cd68* mRNA than males (**Fig. 2b**; sex: p=0.05). In the vHipp, DEP/MS upregulated *Cd11b* mRNA expression (**Fig. 2c**; treatment: p=0.04). There was also a trend towards an interaction effect (interaction: p=0.09). Tukey’s post hoc comparisons found that this effect was mainly driven by increased expression in the males (*post hoc* VEH/CON vs DEP/MS males: p=0.024) rather than the females (p=0.74). No effects of sex or treatment were observed on expression of *Cd68* or *Tlr4* (**Fig. 2c**). *Il1*β did not reliably amplify in this brain region, suggesting lack of expression and thus was not included in analysis. In the LS, we observed effects of sex, but no effects of treatment. *Cd68* and *Tlr4* mRNA were expressed more highly in females than in males (**Fig. 2d**; *CD68* sex: P=0.0176, *TLR4* sex: P=0.0165). There was also a trend towards an increase in *Il1*β expression in females (**Fig. 2d**; sex: p=0.08). In the AMY, there was a trend towards an effect of sex on *Cd68* expression, with females expressing more *Cd68* than males (**Fig. 2e**; sex: p=0.09). No other effects of sex or treatment on gene expression in the AMY were observed (**Fig. 2e**).

**Table 2.** Neuroimmune gene expression result of 2-way ANOVA [sex x treatment]. Significant statistics (p<0.05) and trends (p<0.1) are bolded. Abbreviations: NAc = nucleus accumbens. LS = lateral septum. AMY = amygdala. dHipp = dorsal hippocampus. vHipp = ventral hippocampus. *Cd11b* = cluster of differentiation 11b (integrin alpha M). *Cd68* = cluster of differentiation 68 gene. *Il1β* = interleukin-1 β gene. *Tlr4* = toll-like receptor 4 gene.

| Region | Gene | Treatment (F) | Treatment (p) | Sex (F) | Sex (p) | Interaction (F) | Interaction (p) |
| --- | --- | --- | --- | --- | --- | --- | --- |
| NAc | *Cd11b* | $F_{(1, 40)} = 4.18$ | **p=0.048** | $F_{(1, 40)} = 0.02$ | p=0.88 | $F_{(1, 40)} = 0.32$ | p=0.58 |
| NAc | *Cd68* | $F_{(1, 41)} = 2.17$ | p=0.15 | $F_{(1, 41)} = 0.04$ | p=0.85 | $F_{(1, 41)} = 1.48$ | p=0.23 |
| NAc | *Il1β* | $F_{(1, 28)} = 0.05$ | p=0.82 | $F_{(1, 28)} = 0.06$ | p=0.81 | $F_{(1, 28)} = 0.43$ | p=0.52 |
| NAc | *Tlr4* | $F_{(1, 36)} = 4.54$ | **p=0.04** | $F_{(1, 36)} = 0.5$ | p=0.48 | $F_{(1, 36)} = 0.47$ | p=0.5 |
| LS | *Cd11b* | $F_{(1, 36)} = 0.31$ | p=0.58 | $F_{(1, 36)} = 2.55$ | p=0.12 | $F_{(1, 36)} = 0.5$ | p=0.49 |
| LS | *Cd68* | $F_{(1, 37)} = 0.62$ | p=0.44 | $F_{(1, 37)} = 6.17$ | **p=0.018** | $F_{(1, 37)} = 0.07$ | p=0.8 |
| LS | *Il1β* | $F_{(1, 33)} = 0.02$ | p=0.88 | $F_{(1, 33)} = 3.24$ | **p=0.081** | $F_{(1, 33)} = 0.05$ | p=0.83 |
| LS | *Tlr4* | $F_{(1, 36)} = 0.12$ | p=0.73 | $F_{(1, 36)} = 6.32$ | **p=0.017** | $F_{(1, 36)} = 0.02$ | p=0.9 |
| AMY | *Cd11b* | $F_{(1, 40)} = 0.22$ | p=0.64 | $F_{(1, 40)} = 1.62$ | p=0.21 | $F_{(1, 40)} = 0.34$ | p=0.57 |
| AMY | *Cd68* | $F_{(1, 39)} = 0.94$ | p=0.34 | $F_{(1, 39)} = 3.11$ | **p=0.086** | $F_{(1, 39)} = 1.94$ | p=0.17 |
| AMY | *Il1β* | $F_{(1, 30)} = 0.002$ | p=0.97 | $F_{(1, 30)} = 0.57$ | p=0.46 | $F_{(1, 30)} = 0.06$ | p=0.81 |
| AMY | *Tlr4* | $F_{(1, 37)} = 0.93$ | p=0.34 | $F_{(1, 37)} = 0.24$ | p=0.63 | $F_{(1, 37)} = 0.23$ | p=0.63 |
| dHipp | *Cd11b* | $F_{(1, 41)} = 1.87$ | p=0.18 | $F_{(1, 41)} = 0.31$ | p=0.58 | $F_{(1, 41)} = 0.1$ | p=0.76 |
| dHipp | *Cd68* | $F_{(1, 37)} = 2.67$ | p=0.11 | $F_{(1, 37)} = 3.96$ | **p=0.054** | $F_{(1, 37)} = 1.3$ | p=0.26 |
| dHipp | *Il1β* | $F_{(1, 29)} = 0.24$ | p=0.63 | $F_{(1, 29)} = 1.98$ | p=0.17 | $F_{(1, 29)} = 0.84$ | p=0.37 |
| dHipp | *Tlr4* | $F_{(1, 38)} = 5.08$ | **p=0.03** | $F_{(1, 38)} = 0.06$ | p=0.81 | $F_{(1, 38)} = 0.77$ | p=0.39 |
| vHipp | *Cd11b* | $F_{(1, 33)} = 4.51$ | **p=0.04** | $F_{(1, 33)} = 0.23$ | p=0.63 | $F_{(1, 33)} = 2.99$ | **p=0.093** |
| vHipp | *Cd68* | $F_{(1, 39)} = 0.52$ | p=0.47 | $F_{(1, 39)} = 0.61$ | p=0.44 | $F_{(1, 39)} = 0.04$ | p=0.85 |
| vHipp | *Tlr4* | $F_{(1, 41)} = 1.16$ | p=0.29 | $F_{(1, 41)} = 0.08$ | p=0.78 | $F_{(1, 41)} = 0.49$ | p=0.49 |

**Figure 1:**
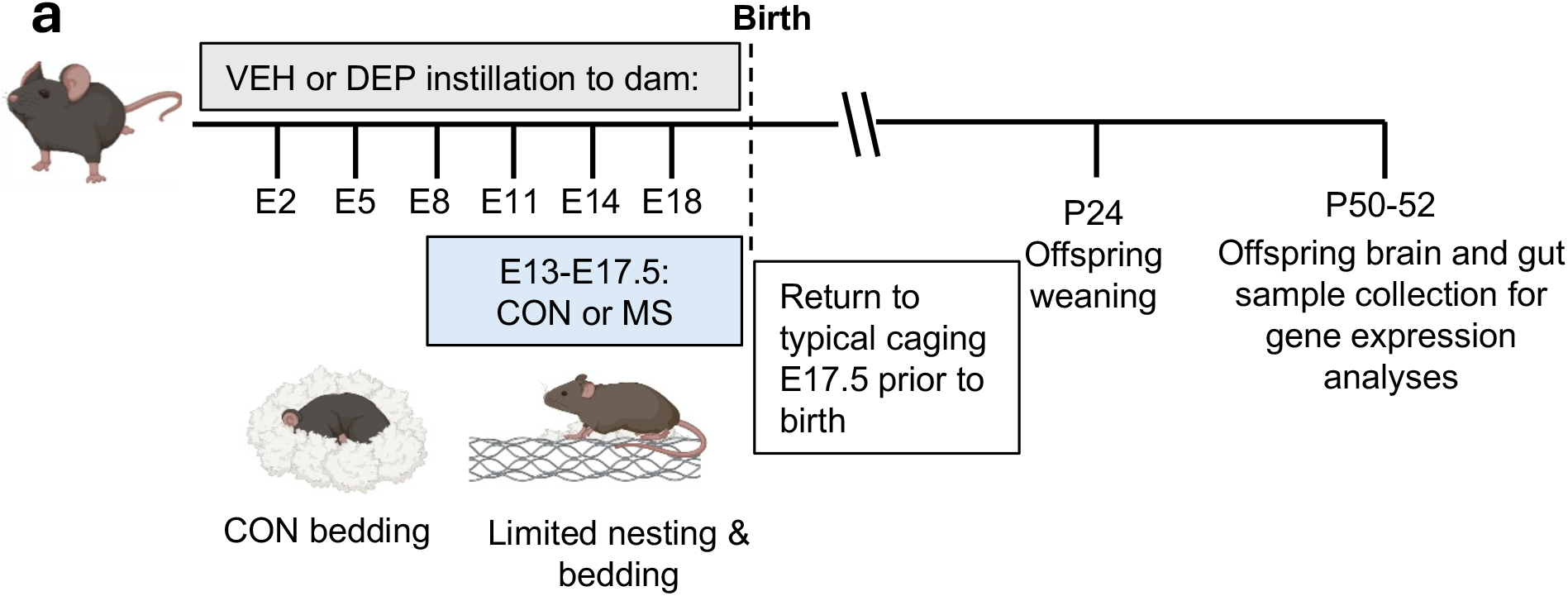
Diagram of the DEP/MS or VEH/CON procedure. Following time-mated pregnancies, dams received either vehicle (VEH) or diesel exhaust particle (DEP) orotracheal instillations every three days throughout pregnancy beginning on embryonic day (E) 2. Between E13-E17.5, dams were housed with either control nesting (CON) containing Alpha-Dri bedding and a full cotton nestlet or maternal stress (MS) nesting containing a wire mesh grate and 2/3 of a cotton nestlet. All exposures were limited to the prenatal period and litters were left undisturbed following birth. Brain and gut tissue samples were collected between P50-52.

**Figure 2:**
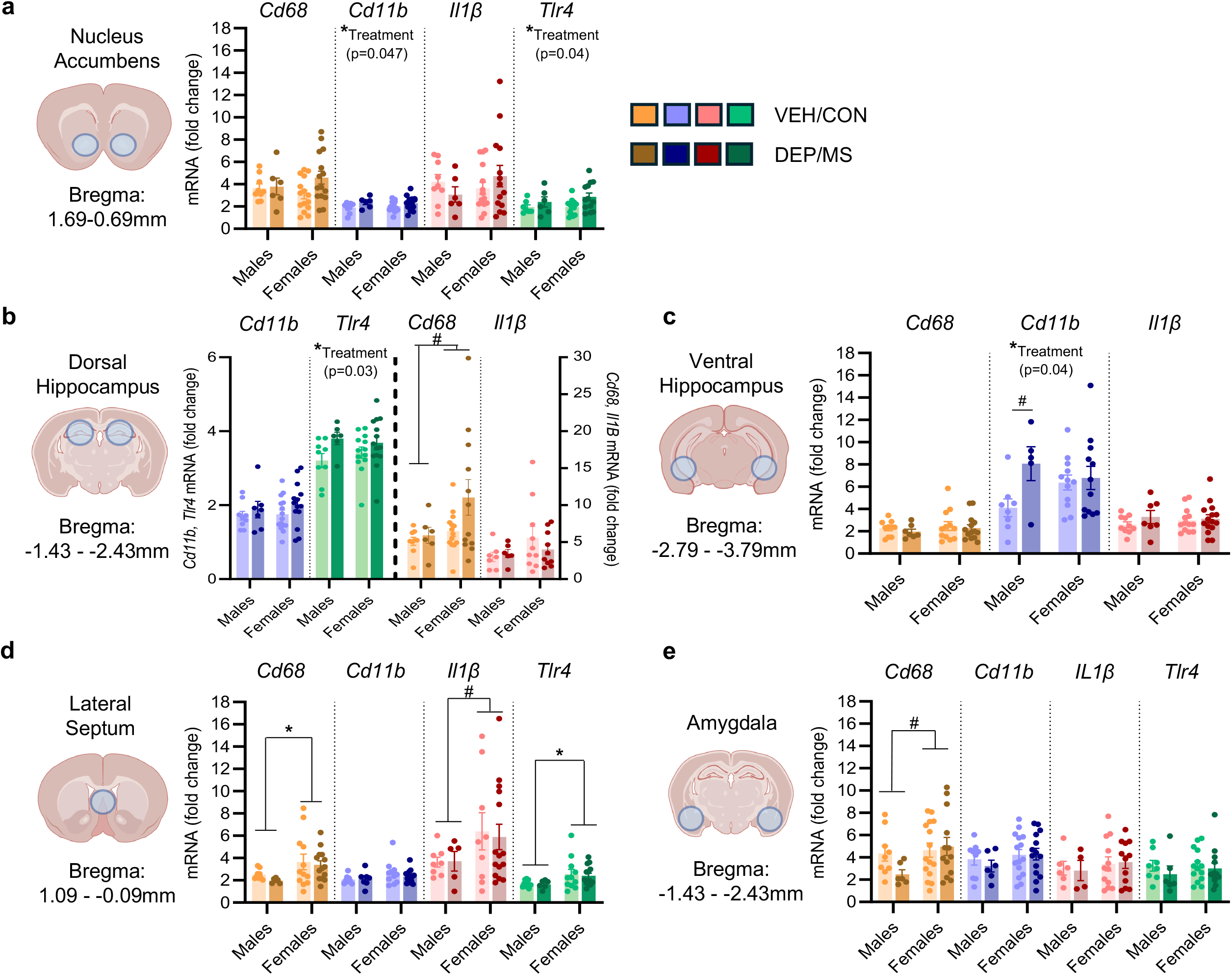
Neuroimmune gene expression in the NAc and hippocampus is impacted by DEP/MS. Gene expression of integrin alpha M subunit of the complement receptor expressed by microglia (*Cd11b*), cluster of differentiation 68 (*Cd68*), interleukin-1β (*Il1*β), and toll-like receptor 4 (*Tlr4*) mRNA in the **(a)** NAc, **(b)** dHipp, **(c)** vHipp **(d)** LS, **(e)** AMY. VEH/CON animals are represented with light colors and DEP/MS with dark colors. Data are represented as mean□+ □SEM, \**p*□< □0.05, #p < 0.1 (2-way ANOVA: sex x treatment).

### 3.2 DEP/MS increases expression of genes related to DA metabolism and reuptake from the synapse in the NAc, but not the hippocampus

In the NAc, while *Drd1* mRNA expression was lower in DEP/MS males as compared to VEH/CON males, this was not statistically significant (**Fig. 3a**, for complete F statistic see **Table 3**). However, we observed a significant interaction effect on *Drd2* mRNA (**Fig. 3a**, p=0.02) whereby DEP/MS significantly decreased expression of *Drd2* in males only (*post hoc* VEH/CON vs. DEP/MS males; p=0.06). DEP/MS increased expression of *Comt* in both sexes (Fig 3a; treatment: p=0.04), and robustly increased expression of *Slc6a3* (**Fig. 3a**; treatment: p<0.001). In the dHipp, there were no effects of sex or treatment on *Drd1, Drd2, Comt*, or *Slc6a3* mRNA expression (**Fig. 3b**). Finally, in the vHipp, DEP/MS decreased *Comt* mRNA expression as compared to control (**Fig. 3c**; p=0.03), opposite of the effect seen in the NAc. No sex or treatment effects were found on *Drd1, Drd2*, or *Slc6a3* mRNA expression.

**Table 3.**
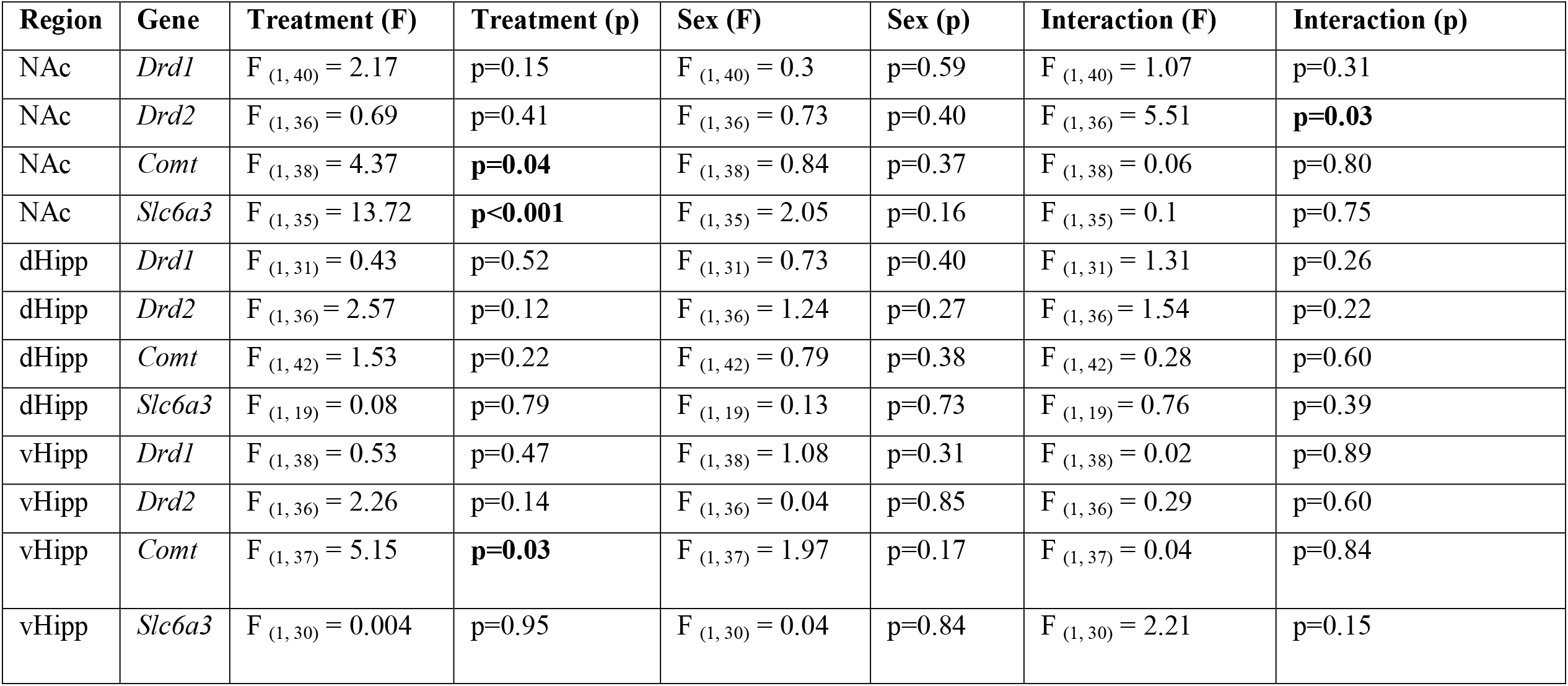
Dopaminergic gene expression results 2-way ANOVA [sex x treatment]. Significant statistics (p<0.05) are bolded. Abbreviations: NAc = nucleus accumbens. dHipp = dorsal hippocampus. vHipp = ventral hippocampus. *Drd1* = dopamine receptor 1 gene. *Drd2* = dopamine receptor 2 gene. *Comt* = catechol-O-methyltransferase gene. *Slc6a3* = dopamine transporter gene.

**Table 4.** Intestinal immune gene expression results 2-way ANOVA [sex x treatment]. Significant statistics (p<0.05) and trends (p<0.10) are bolded. Abbreviations: *Il1β* = interleukin-1 β gene. *Il17ra* = interleukin-17 receptor A gene. *Tlr4* = toll-like receptor 4 gene. *Tnf-α* = tumor necrosis factor alpha gene.

| Region | Gene | Treatment (F) | Treatment (p) | Sex (F) | Sex (p) | Interaction (F) | Interaction (p) |
| --- | --- | --- | --- | --- | --- | --- | --- |
| Jejunum | *Il1β* | F<sub>(1, 16)</sub> = 0.73 | p=0.40 | F<sub>(1, 16)</sub> = 4.87 | **p=0.04** | F<sub>(1, 16)</sub> = 1.077e-006 | p=0.99 |
| Jejunum | *Il17ra* | F<sub>(1, 22)</sub> = 0.65 | p=0.43 | F<sub>(1, 22)</sub> = 1.36 | p=0.25 | F<sub>(1, 22)</sub> = 0.17 | p=0.68 |
| Jejunum | *Tlr4* | F<sub>(1, 20)</sub> = 3.88 | **p=0.06** | F<sub>(1, 20)</sub> = 10.85 | **p<0.01** | F<sub>(1, 20)</sub> = 5.54 | **p=0.03** |
| Jejunum | *Tnf-α* | F<sub>(1, 20)</sub> = 0.05 | p=0.82 | F<sub>(1, 20)</sub> = 3.56 | **p=0.07** | F<sub>(1, 20)</sub> = 2.35 | p=0.14 |
| Ileum | *Il1β* | F<sub>(1, 38)</sub> = 0.0002 | p=0.99 | F<sub>(1, 38)</sub> = 3.39 | **p=0.07** | F<sub>(1, 38)</sub> = 1.45 | p=0.24 |
| Ileum | *Il17ra* | F<sub>(1, 34)</sub> = 0.12 | p=0.74 | F<sub>(1, 34)</sub> = 3.28 | **p=0.08** | F<sub>(1, 34)</sub> = 0.87 | p=0.36 |
| Ileum | *Tlr4* | F<sub>(1, 40)</sub> = 0.85 | p=0.36 | F<sub>(1, 40)</sub> = 1.35 | p=0.25 | F<sub>(1, 40)</sub> = 0.22 | p=0.65 |
| Ileum | *Tnf-α* | F<sub>(1, 31)</sub> = 0.17 | p=0.69 | F<sub>(1, 31)</sub> = 6.19 | **p=0.02** | F<sub>(1, 31)</sub> = 0.5 | p=0.48 |
| Colon | *Il1β* | F<sub>(1, 19)</sub> = 1.7 | p=0.21 | F<sub>(1, 19)</sub> = 0.01 | p=0.91 | F<sub>(1, 19)</sub> = 2.63 | p=0.12 |
| Colon | *Il17ra* | F<sub>(1, 24)</sub> = 3.56 | **p=0.07** | F<sub>(1, 24)</sub> = 2.52 | p=0.13 | F<sub>(1, 24)</sub> = 8.72 | **P<0.01** |
| Colon | *Tlr4* | F<sub>(1, 22)</sub> = 1.84 | p=0.19 | F<sub>(1, 22)</sub> = 0.02 | p=0.90 | F<sub>(1, 22)</sub> = 5.51 | **p=0.03** |
| Colon | *Tnf-α* | F<sub>(1, 25)</sub> = 1.09 | p=0.31 | F<sub>(1, 25)</sub> = 5.16 | **p=0.03** | F<sub>(1, 25)</sub> = 1.85 | p=0.19 |

**Figure 3:**
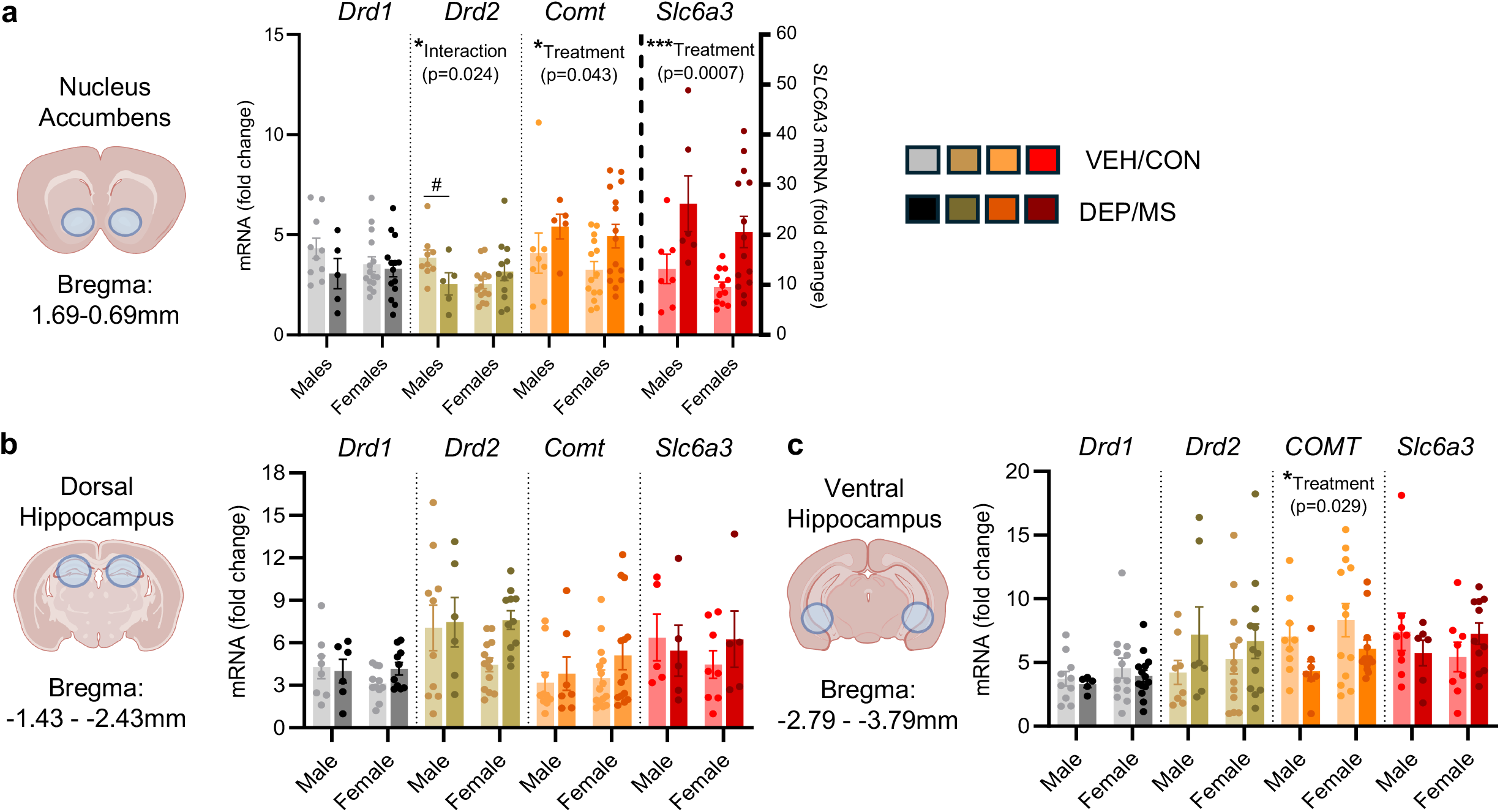
DEP/MS alters expression of genes related to DA metabolism and reuptake from the synapse in the NAc and vHipp and has sex-specific effects on *Drd2* expression. Gene expression of *Drd1, Drd2*, catechol-O-methyltransferase (*Comt*), and dopamine transporter (*Slc6a3*) mRNA in the **(a)** NAc, **(b)** dHipp, **(c)** vHipp **(d)** LS, **(e)** AMY. VEH/CON animals are represented with light colors and DEP/MS with dark colors. Data are represented as mean□+□SEM, \**p*□<□0.05, ***p < 0.001 (2-way ANOVA: sex x treatment). In the case of significant interaction effects, Tukey’s *post hoc* comparisons are labeled.

### 3.3 DEP/MS decreases expression of peripheral immune-related genes in males and increases them in females in the small and large intestine

We assessed gene expression in two regions of the small intestine, the jejunum and the ileum, as well as in the large intestine (colon; see **Fig. 4a** for schematic). In the jejunum, we observed sex differences in 3 of the 4 genes expressed. Specifically, females had higher expression of *Il1*β and *Tlr4* mRNA than males, but a trend towards lower *Tnfα* (**Fig. 4b**, main effects of sex: *Il1*β: sex: p=0.04, *Tlr4* sex: p<0.01, *Tnfα* sex: p=0.07). There was also a trend towards a treatment effect and a significant interaction effect on *Tlr4* mRNA expression such that DEP/MS increased *Tlr4* mRNA in females (**Fig. 4b**, treatment p=0.06; interaction p=0.02; *post hoc* VEH/CON vs. DEP/MS females; p=0.02). We also observed higher immune gene expression in females in the ileum (**Fig. 4c**). Females expressed more *Tnfα* mRNA than males (**Fig. 4c**; sex: p=0.02) and trended towards expressing more *Il1*β and *Il17ra* mRNA than males (**Fig. 4c**; *Il1*β sex: p=0.07, *Il17ra* sex: p=0.08). There were no significant effects of treatment or sex on *Tlr4* mRNA expression in the ileum (**Fig. 4c**). Of the intestinal regions tested, DEP/MS had the greatest effects on gene expression in the colon (**Fig. 4d**). There were significant interaction effects between sex and treatment on *Il17ra* and *Tlr4 mRNA* expression, whereby DEP/MS exposure increased expression in females but not males (**Fig. 4d**; *Il17ra* treatment: p=0.07, interaction: p=0.007, *post hoc* VEH/CON vs. DEP/MS females p=0.01; *Tlr4* interaction: p=0.03, *post hoc* VEH/CON vs. DEP/MS females p=0.07). Females expressed more *Tnfα* mRNA in the colon than males (**Fig. 4d**; sex: p=0.03). Finally, there were no effects of sex or treatment on *Il1*β expression in the colon (**Fig. 4d**).

**Figure 4:**
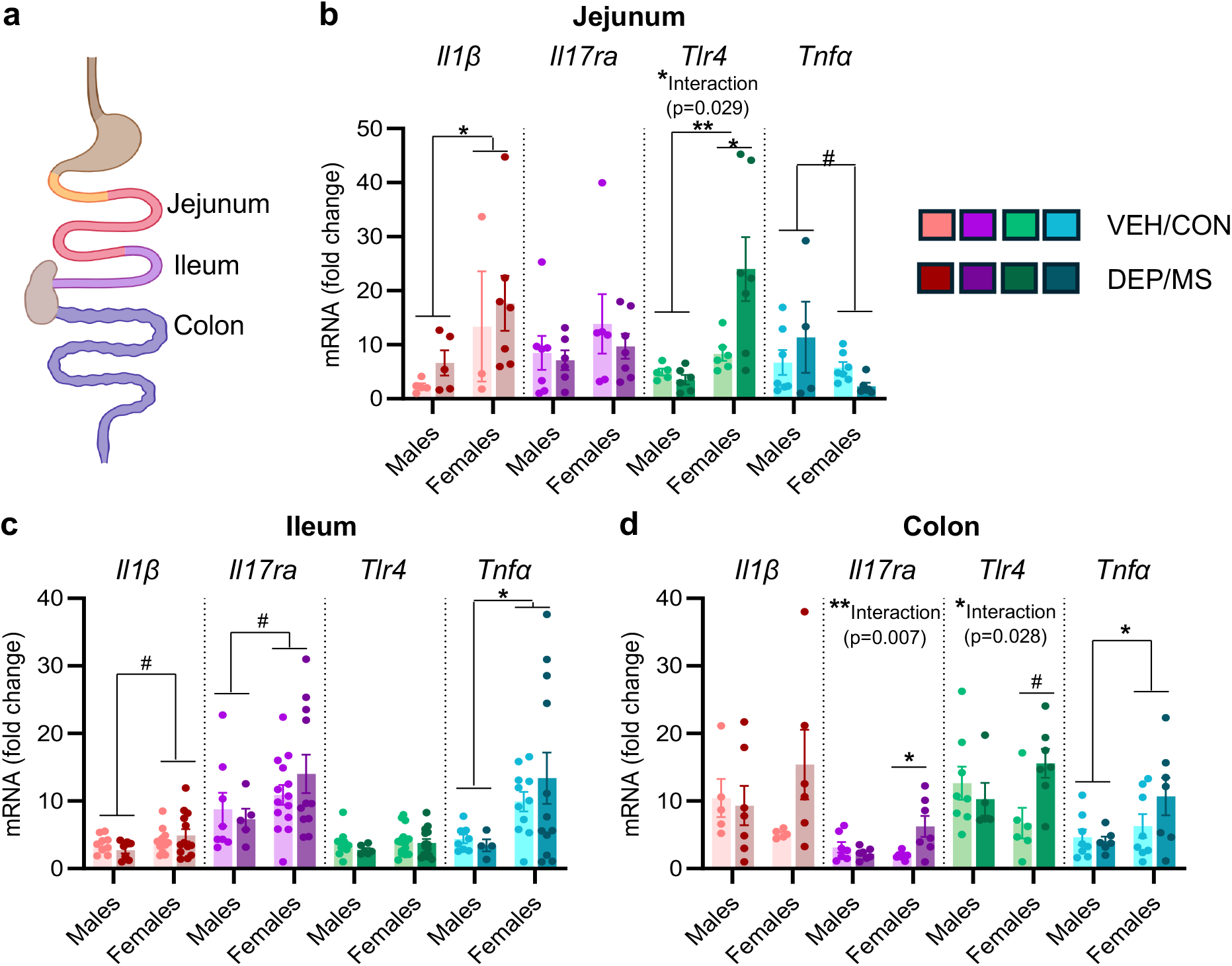
DEP/MS increases expression of peripheral immune-related genes in females in the small and large intestine. **(a)** Diagram of the mouse small and large intestine. **(b-d)** Gene expression of interleukin-1β (*Il1*β), interleukin-17 receptor *(Il17ra)*, toll-like receptor 4 (*Tlr4*), and tumor necrosis factor alpha (*Tnf-a)* mRNA in the **(b)** jejunum, **(c)** ileum, and **(d)** colon. VEH/CON animals are represented with light colors and DEP/MS with dark colors. Data are represented as mean□+□SEM, #p < 0.1, \**p*□<□0.05, **p < 0.01 (2-way ANOVA: sex x treatment). In the case of significant interaction effects, Tukey’s *post hoc* comparisons are labeled.

## 2. DISCUSSION

Our results suggest several key findings. First, *CD11b* and *Tlr4* were the most-altered immune genes following DEP/MS exposure, with significant treatment effects in the NAc, dHipp, and vHipp, but not in the LS or AMY. Second, we observed robust changes in dopaminergic gene expression in the NAc following DEP/MS, consistent with a phenotype of reduced dopamine signaling and increased reuptake and catabolism. Third, immune gene expression changes in the intestine following DEP/MS were female-specific: *Tlr4* mRNA was increased following DEP/MS in females in the jejunum and colon and *IL-17ra* mRNA was increased in the colon. Finally, we observed numerous sex differences in immune gene expression regardless of treatment, typically with higher expression in females, in brain regions with known sex differences in function (such as the LS and AMY) as well as in all regions of the intestines, but most commonly the ileum. Overall, these findings highlight the NAc as a key site of neuroimmune and dopaminergic changes following DEP/MS exposure in both sexes and reveal female-specific increases in immune gene expression in the intestine following DEP/MS exposure.

The brain region in which we observed the most robust changes in gene expression following DEP/MS exposure was the NAc. We found that NAc *CD11b* and *Tlr4* mRNA expression were increased following DEP/MS as compared to control regardless of sex. Furthermore, we found that *Drd2* expression was decreased in males but not females in the NAc, while *Comt* and *Slc6a3* were increased in both sexes following DEP/MS as compared to VEH/CON. These findings are in line with our previous work in this model suggesting that the NAc is an important site for neuroimmune and dopaminergic changes following DEP/MS exposure. Specifically, we previously reported microglial hyper-ramification, increased microglial volume, and differential microglia gene expression in the NAc of adolescent males but not females following DEP/MS exposure (Smith et al., 2023a). Furthermore, we found that DEP/MS exposure decreased NAc *Drd1* and *Drd2* mRNA expression in males but not females and that chemogenetic activation of dopaminergic neurons in the ventral tegmental area was sufficient to rescue social behavior deficits in male DEP/MS-exposed offspring (Smith et al., 2023a). Our findings here recapitulate the sex-specific effects of DEP/MS on *Drd2* mRNA expression that we had previously observed (as well as the pattern for *Drd1* expression although the differences did not reach significance here) and extend them by showing that *Comt* and *Slc6a3* are also impacted by DEP/MS in both sexes.

The *Comt* gene encodes for catecholamine-O-methyltransferase, a key enzyme in catecholamine catabolism pathways. For example, *Comt* converts 3,4-dihydroxyphenylacetic acid (DOPAC, a main metabolite of dopamine) into homovanillic acid (HVA). *Slc6a3* encodes for dopamine transporter 1 (DAT1) which is responsive for dopamine reuptake into the presynapse. Together, the heightened expression of both these molecules in the NAc that we observe following DEP/MS exposure suggests increased dopamine degradation and reuptake at the synapse. One important caveat, however, is that these findings are only at the gene expression level and we do not have a concurrent measure of dopamine level itself. Therefore, it is also possible that this difference could reflect higher starting dopamine content in the NAc. This may be unlikely given that we previously found a strong trend towards lower tyrosine hydroxylase immunoreactivity in the NAc, at least in males, following DEP/MS (Smith et al., 2023b). Still, it would be of interest to characterize this further, as previous work investigating postnatal exposure to various air pollutants has found increased protein content for dopamine and its’ metabolites in the striatum. For example, exposure to inhaled copper, a common component of air pollution, increases striatal dopamine (DA), DOPAC, and HVA as compared to control, along with a decrease in dopamine turnover (HVA/DA) in males at P14 (Cubello et al., 2024). Similarly, postnatal exposure to combined iron and sulfur dioxide inhalation increased striatal DA, DOPAC, and HVA in male but not female mice (Sobolewski et al., 2018). In rats, Yakota et al. (2016) observed increased NAc-DA in socially isolated males following prenatal DEP exposure as compared to control (Yokota et al., 2016). These findings suggest that the dopamine system is highly sensitive to air pollutant exposures but that the direction of toxicant effects is variable.

Our findings of increased *CD11b* and *Tlr4* mRNA expression in the NAc and hippocampus are in line with broader work suggesting that *CD11b* and *Tlr4* mRNA are altered by developmental air pollution exposure in the brain and body. For instance, in whole brain homogenate, *Tlr4* mRNA is increased in males but not females at P30 following DEP/MS exposure but not following DEP or MS alone (Bolton et al., 2013). Similarly, Bolton et al. (2014) found that *Cd11b* and *Tlr4* mRNA expression increased in both males and females in adipose tissue following exposure to DEP during gestation and high-fat diet during the postnatal period (Bolton et al., 2014). TLR4 is a critical pattern recognition receptor for both pathogen- and danger associated molecular patterns. Interestingly, prior evidence suggests that TLR4 signaling may be a critical mechanism by which microglia respond to DEP exposure. Bolton et al. (2017) found that exposure to DEP alone during gestation increased cortical and hippocampal volumes and increased the proportion of ‘round’ and ‘stout’ microglia in the fetal brain of male offspring. Importantly, these increases were absent in whole-body TLR4 knock out mice (Bolton et al., 2017). In vitro work also supports the idea that TLR4 signaling is necessary for at least some of the downstream effects of air pollution in the brain. Woodward et al. (2017) found that in mixed-glial cultures (microglia and astrocytes), nanoscale particulate matter exposure increased the expression of several proinflammatory cytokines such as TNFα, Il-1β, and Il-6, and these increases were attenuated by treatment with TLR4 siRNA(Woodward et al., 2017). In cultured human microglial cells (line 3; HMC3), PM_2.5_ exposure increases Il-6 and COX2 gene expression and this was prevented by administration of a TLR4 neutralizing antibody (Zhang et al., 2024). Future work should aim to determine the necessity of TLR4 activation to the effects of DEP/MS on dopaminergic gene expression observed here.

We began this study with the hypothesis that the effects of DEP/MS exposure might extend beyond the NAc to other brain regions that are critical to social behavior. We observed treatment effects in the hippocampus (increased *Tlr4* expression in the dHipp and increased *CD11b*/decreased *Comt* in the vHipp) but no treatment effects in the LS or AMY, suggesting that this is only partially the case. Interestingly, while we did not observe effects of DEP/MS on gene expression in these regions, we did find sex differences in gene expression. For example, in the LS, *CD68* mRNA expression was higher in females than in males and *Il1*β trended towards higher expression in females as did *CD68* mRNA in the AMY. CD68 is a lysosomal marker that is expressed almost exclusively in microglia in the central nervous system. Therefore, these findings might suggest higher microglial phagocytosis in females in these brain regions. Our work has important limitations in this context. First, our investigation is limited to the analysis of gene expression, so it is possible that these findings do not translate to the protein level. Second, the use of tissue punches to collect samples means that it is impossible to delineate sub-nuclei in brain regions like the AMY where there are many distinct subregions with important functional differences. Future studies should aim to further explore these findings using methods such as immunohistochemistry that allow for cell type- and sub-region-specificity.

In addition to investigating cytokine expression in the brain, we assessed cytokine gene expression in both the small and large intestine. While most of the treatment effects that we observed in the brain did not depend on sex, all the effects of DEP/MS on intestinal gene expression were female specific. We also observed several baseline sex differences in immune gene expression in the gut. We found that *Il1*β expression was higher or tended to be higher in females than males in the jejunum and ileum respectively. *Tnfα* expression trended towards being lower in the jejunum of females as compared to males and was significantly lower in both the ileum and colon. DEP/MS exposure increased *Tlr4* in the jejunum and colon in females only, and increased *Il17ra* in females in the colon. *Il17ra* also trended towards being higher in females than in males in the ileum. The sex differences we observe are in line with previous investigations suggesting sex differences in mucosal immunity. For example, in humans, IL-1β, TNF, and IL-17 proteins are all higher in jejunum mucosal biopsies from women as compared to men (Sankaran-Walters et al., 2013). In contrast, the female-specific nature of our effects appears to be opposite to findings in many models of intestinal inflammation such as colitis, high-fat diet, and even air pollution in which males tend to exhibit more changes in intestinal inflammation than females (Brettle et al., 2022; Fitch et al., 2020; Guilloteau et al., 2022; Hases et al., 2023). Indeed, estrogen receptor signaling in the intestine may protect females from the inflammatory consequences of DSS-induced colitis relative to males (Pereda et al., 2025). However, Fitch et al. (2020) did find increased colonic Tlr4 expression following woodsmoke inhalation in adulthood, albeit in males (Fitch et al., 2020). It remains to be determined what the functional consequences of these sex-specific effects of DEP/MS might be.

## Conclusions and Future Directions

Our results suggest several interesting avenues for future investigation. First, we observed the most robust changes in dopaminergic and immune gene expression in the NAc. Future studies should aim to determine whether dopamine release is blunted in the NAc following DEP/MS exposure and what the functional consequences of this for behavior might be. Moreover, given that changes in NAc-DA coincide with changes in immune markers in the NAc, it would be of interest to determine whether and how neuroimmuity shapes dopaminergic function in this context. We also found female-specific changes in immunity in the gut following DEP/MS exposure. It remains unclear what the functional significance is of elevated immune gene expression in females following DEP/MS exposure and whether this is protective or harmful. Together, these findings expand our understanding of the impact of prenatal air pollution and maternal stress on immune and dopaminergic gene expression in the offspring gut-brain axis.

## Acknowledgements

We would like to thank the animal care staff at Boston College for their help and care of our animals and all the members of the Smith Lab for their feedback on the manuscript.

## CRediT Authorship Contribution Statement

Caroline J. Smith: funding acquisition, conceptualization, writing - review and editing, methodology, project administration.

Elise M. Martin: conceptualization, investigation, writing - original draft, writing - review and editing, formal analysis, visualization, supervision.

Matthew J. Morales: writing - original draft, writing - review and editing, visualization.

Niki Y. Li: conceptualization, investigation, formal analysis.

Maura C. Stoehr: conceptualization, investigation.

Matthew J. Kern: investigation, formal analysis.

Madeline F. Winters: investigation, formal analysis.

